# Adaptive responding to stimulus-outcome associations requires noradrenergic transmission in the medial prefrontal cortex

**DOI:** 10.1101/2023.12.22.572995

**Authors:** Alessandro Piccin, Hadrien Plat, Mathieu Wolff, Etienne Coutureau

**Affiliations:** Université de Bordeaux, INCIA, UMR 5287, 33000 Bordeaux, France; CNRS, INCIA, UMR 5287, 33000 Bordeaux, France

## Abstract

A dynamic environment, such as the one we inhabit, requires organisms to continuously update their knowledge of the setting. While the prefrontal cortex is recognized for its pivotal role in regulating such adaptive behavior, the specific contributions of each prefrontal area remain elusive. In the current work, we investigated the direct involvement of two major prefrontal subregions, the medial prefrontal cortex (mPFC) and the ventrolateral orbitofrontal cortex (vlOFC), in updating Pavlovian stimulus-outcome (S-O) associations following contingency degradation. Specifically, animals had to learn that a specific cue, previously fully predicting the delivery of a specific reward, was no longer a reliable predictor. First, we found that chemogenetic inhibition of mPFC, but not of vlOFC, neurons altered the rats’ ability to adaptively respond to degraded and non-degraded cues. Next, given the growing evidence pointing at noradrenaline (NA) as a main neuromodulator of adaptive behavior, we decided to investigate the possible involvement of NA projections to the two subregions in this higher-order cognitive process. Employing a pair of novel retrograde vectors, we traced NA projections from the locus coeruleus (LC) to both structures and observed an equivalent yet relatively segregated amount of inputs. Then, we showed that chemogenetic inhibition of NA projections to the mPFC, but not to the vlOFC, also impaired the rats’ ability to adaptively respond to the degradation procedure. Altogether, our findings provide important evidence of functional parcellation within the prefrontal cortex and point at mPFC-NA as key for updating Pavlovian S-O associations.

**Significant statement:** The ability to update stimulus-outcome (S-O) associations is a key adaptive behavior, essential for surviving in an ever-changing environment. The prefrontal cortex is well-known for playing a key role in this process. The discrete contribution of each prefrontal subregion and of different neurotransmitters, however, remains unclear. In the current study, we show that inhibiting medial prefrontal (mPFC), but not ventrolateral orbitofrontal cortex (vlOFC), neurons impairs the rats’ ability to update S-O associations following contingency degradation. Moreover, we demonstrate that discrete noradrenergic projections to the two subregions exist and that inhibiting the ones projecting to the mPFC, but not to the vlOFC, once again impairs the rats’ behavior, thereby implying a substantial contribution of noradrenaline in orchestrating this higher-order cognitive process.

## Introduction

The ability to learn from experience is a key adaptive behavior, essential for surviving and thriving in a constantly changing environment. Perhaps not surprisingly, this ability is hampered in several neuropsychiatric conditions, ranging from developmental to neurocognitive disorders (Uddin, 2021). Investigating the neural mechanisms that underlie this higher-level cognitive process is therefore crucial to advance our understanding of these debilitating diseases.

Among all brain areas, the prefrontal cortex is well-known for playing a pivotal role in adaptive behavior, specifically in the encoding and updating of instrumental action-outcome (A-O) and of Pavlovian stimulus-outcome (S-O) associations, with a high parcellation of function existing among prefrontal subregions. For instance, the medial prefrontal cortex (mPFC) has been historically linked to the acquisition (Killcross and Coutureau, 2003; Tran-Tu-Yen et al., 2009) and updating (Boitard et al., 2016) of instrumental associations, and more recently has been suggested to participate in the updating of Pavlovian associations following outcome devaluation (Niedringhaus and West, 2022), while the orbitofrontal cortex (OFC) has been implicated, directly or indirectly, in the updating of previously established instrumental (Parkes et al., 2018; Fresno et al., 2019; Cerpa et al., 2023) and Pavlovian associations (Ostlund and Balleine, 2007; Alcaraz et al., 2015; Morceau et al., 2022).

The neuromodulation of such adaptive behaviors has long been associated with dopamine (DA), from the notion that associative learning is driven by DA transients that correlate with prediction errors (Schultz, 2016; Sharpe et al., 2017; Langdon et al., 2018), to the fact that optogenetic stimulation of DA neurons paired with a cue has been shown to evoke Pavlovian conditioned responses (Saunders et al., 2018). However, recent theoretical models (Cerpa et al., 2021) propose that it might be crucial to take into account the other main catecholaminergic neurotransmitter, noradrenaline (NA), a key regulator of attention and arousal in the central nervous system, which is believed to track uncertainty and drive adaptive behavior (Bouret and Sara, 2004; Tait et al., 2007; McGaughy et al., 2008; Tervo et al., 2014; Uematsu et al., 2017; Jahn et al., 2018; Cope et al., 2019). Notably, the mPFC and the OFC are believed to receive discrete and direct far-reaching NA projections from the brainstem locus coeruleus (LC), the primary source of NA to the mammalian cortex (Agster et al., 2013; Chandler et al., 2014, 2013). This notion has been recently backed up by the fact that chemogenetic inhibition of LC-NA projections to the ventral and lateral parts of the OFC (vlOFC), but not to the mPFC, impaired the rats’ ability to update A-O associations following reversal (Cerpa et al., 2023). To our knowledge, this specific anatomical dissociation of effects (mPFC vs vlOFC) and the possible implication of NA in the updating of S-O associations have never been investigated.

Therefore, in the current work, we looked into the discrete involvement of these two major prefrontal subregions in Pavlovian contingency degradation, a task in which rats are challenged to learn that a cue, previously the sole predictor of a specific reward, is no longer a better predictor than the absence of that same cue. First, we found that chemogenetic inhibition of mPFC, but not vlOFC, neurons during the degradation phase altered the rats’ ability to adaptively respond to degraded and non-degraded cues. Next, employing a pair of novel retrograde vectors, we traced LC-NA projections to both structures and observed an equivalent yet relatively segregated amount of inputs. At last, we showed that chemogenetic inhibition of NA projections to the mPFC, but not to the vlOFC, once again impaired the rats’ ability to adapt to the degradation of S-O contingencies.

The present work contributes to integrate and revisit our knowledge of how the prefrontal cortex controls flexible, goal-directed behaviors. Specifically, we question the implication of the OFC in updating S-O contingencies, and point at the mPFC and its NA innervation instead as crucial for executing such behavioral adaptation.

## Materials and methods

### Subjects and housing conditions

A total of 95 male Long-Evans rats, aged 2-3 months, were acquired from the Centre d’Elevage Janvier (France). Animals were housed in pairs with *ad libitum* access to water and standard lab chow, then put on food restriction 2 days before the start of behavioural experiments and maintained at approximately 90% of their *ad libitum* feeding weight. Rats were handled daily for 3 days before the beginning of the experiments. The facility was maintained at 21 ± 1 °C on a 12 hr light/dark cycle (lights on at 8:00 am). Environmental enrichment was provided by tinted polycarbonate tubing elements, in accordance with current French (Council directive 2013-118, February 1, 2013) and European (directive 2010-63, September 22, 2010, European Community) laws and policies regarding animal experimentation. The experiments received approval from the ethics and animal experimentation committee of the French Ministry of Higher Education and Innovation (reference number of the project: APAFIS#27928-2020110918011853 v2).

### Viral vectors

In Experiment 1, an adeno-associated viral vector carrying the inhibitory hM4Di designer receptor exclusively activated by designer drugs (DREADDs) was acquired from the Viral Vector Production Unit at Universitat Autonoma de Barcelona, Spain. This AAV8-CaMKII-hM4D(Gi)-mCherry vector was derived from a plasmid obtained from Addgene (pAAV-CaMKIIa-hM4D(Gi)-mCherry; Addgene plasmid #50477) and used at a titer of 3.5 x 10^12 GC/mL, as validated in a previous study (Parkes et al., 2018). In Experiment 2, we employed two retrograde canine adenovirus type 2 (CAV-2) vectors harboring a PRS promoter that enables the expression of fluorescent proteins in noradrenergic neurons projecting to the target regions of interest. Both CAV-2-PRS-FsRed and CAV-2-PRS-eGFP were supplied by Marina Lavigne and Eric J. Kremer at the Institut Genetique Moleculaire, CNRS UMR 5535, Montpellier, France. They were used at a concentration of 3.6 x 10^12 VP/mL and 3.45 x 10^12 VP/mL, respectively. In Experiment 3, we employed another CAV-2 vector equipped with a PRS promoter, this time controlling the expression of an hM4Di receptor and therefore enabling selective chemogenetic targeting of noradrenergic projections. This CAV-2-PRS-HA-hM4Di-hSyn-mCherry vector was also generously provided by Eric Kremer’s team and used at a titer of 3.5 x 10^12 VP/mL, as validated in a previous study (Cerpa et al., 2023).

### Stereotaxic surgery

For all experiments, rats were anesthetized with 5% inhalant isoflurane gas with oxygen and placed in a stereotaxic frame with atraumatic ear bars (Kopf Instruments) in a flat skull position. Anesthesia was maintained with 1.5% isoflurane and complemented with a subcutaneous injection of ropivacaïne (a bolus of 0.15 mL at 2 mg/mL) at the incision site and then disinfected using betadine. All viral vectors were injected using a microinjector (UMP3 UltraMicroPump II with Micro4 Controller, World Precision Instruments) connected to a 10 µL Hamilton syringe. For Experiment 1 and Experiment 3, each animal received two injections per hemisphere of either AAV8-CaMKII-hM4D(Gi)-mCherry or CAV-2-PRS-HA-hM4Di-hSyn-mCherry viruses in the mPFC or in the vlOFC (VO+LO). For Experiment 2, animals received counterbalanced unilateral injections of CAV-2-PRS-eGFP and CAV-2-PRS-FsRed in the mPFC and the vlOFC (VO/LO). Each injection consisted of 1 µL infused at a rate of 200 nL/min. After each injection, the syringe remained in place for an additional 10 min before being removed. The coordinates used for the vlOFC were the following: AP +3.7, ML ± 2.0, DV −5.0 and AP +3.2, ML ± 2.8, DV −5.2 for both AAV8 and CAV-2 injections. The coordinates used for mPFC injections were the following: AP +3.2, ML ± 0.6, DV −3.4 and AP +3.0, ML ± 0.7, DV −5.4 or AP +3.8, ML ± 0.6, DV −3.6 and AP +3.2, ML ± 0.6, DV −3.6 for AAV8 or CAV-2 injections, respectively. During surgery, a heating pad was placed under each rat to maintain its body temperature. Following surgery, rats were subcutaneously administered with a nonsteroidal anti-inflammatory drug (meloxicam, 2 mg/mL/kg) and individually housed in a warmed cage with facilitated access to food and water for 2 hours post-surgery. Rats underwent a recovery period of 3 weeks to allow proper transgene expression. Injection sites were confirmed histologically after the completion of behavioural experiments.

### Chemogenetics

The DREADDs agonist deschloroclozapine (DCZ, MedChemExpress) was dissolved in dimethyl sulfoxide (DMSO) at a 50 mg/mL concentration and stored at −80°C (stock solution). The stock solution was then diluted in sterile saline to a final concentration of 0.1 mg/kg and administered intraperitoneally (i.p., 10 mL/kg) 40-45 min before behavioral experiments, as validated in a previous study (Cerpa et al., 2023). The vehicle solution (Veh) was made using 0.2% DMSO in sterile saline. Injectable solutions were freshly prepared on the day of use and manipulated under low-light conditions.

### Behavioural apparatus

Training and testing procedures took place in 8 identical operant chambers (40 cm width x 30 cm depth x 35 cm height, Imetronic, Pessac, France) individually enclosed in sound and light-resistant wooden chambers (74 x 46 x 50 cm). Each chamber was equipped with 2 pellet dispensers that delivered grain (Rodent Grain-Based Diet, 45mg, Bio-Serv) or sugar (LabTab Sucrose Tablet, 45 mg, TestDiet) pellets into a food port (magazine) when activated. Two retracted levers were located on each side of the magazine. Each chamber had a ventilation fan producing a background noise of 55 dB and was illuminated by four LEDs in the ceiling. Experimental events were controlled and recorded by a computer located in the room and equipped with the POLY software (Imetronic, Pessac, France). Rats were habituated to the magazine through 3 daily sessions of training before the beginning of the experiments. During these sessions, they received 40 sugar and 40 grain pellets in the magazine, pseudo-randomly interspersed, on average every 60 seconds.

### Pavlovian conditioning

Rats first underwent 8 x 40 min daily sessions of Pavlovian conditioning, during which 2 auditory CS (tone and clicker) were paired with 2 distinct food rewards (sucrose and grain). Animals were divided into 2 groups, one with tone-sucrose and clicker-pellet pairings, and the other with the opposite combinations. Each CS was presented for 20 sec, during which the corresponding food reward was delivered twice in a pseudo-random manner. Auditory cues were presented each day in 2 blocks of 15 CS with alternate starting order. Rats therefore received a total of 60 rewards per daily session. The time between each CS was pseudo-randomized and ranged from 30 to 90 sec, with an average of 60 sec.

### Pavlovian contingency degradation

Following Pavlovian conditioning, rats experienced 6 daily sessions of contingency degradation during which the predictability of one of the cues was degraded with non-contingent delivery of the reward during the interval period of the CS (60 sec on average). Specifically, the 30 reward deliveries occurred at the same frequency both during and outside of the CS presentation. The degraded CS was therefore no longer a reliable predictive cue of the reward. On the other hand, the non-degraded CS was kept at the same schedule of the Pavlovian conditioning, with 2 rewards being delivered during each 20 sec CS, with pseudo-randomized 60 sec average intervals (± 30 sec). During the degradation sessions, rats were injected systemically (i.p., 10 mL/kg) with either DCZ (0.1 mg/kg) or Veh solutions 40-45 min prior being placed in the operant chambers. All CS–O associations and schedules (Degraded vs Non-degraded) were counterbalanced between experimental groups.

### Test in extinction

Following the last session of contingency degradation, rats underwent 2 tests in extinction, according to a within-subject experimental design. Specifically, each experimental group was divided into 2 subgroups, one receiving DCZ during the first test, and the other receiving Veh. Conditions were reversed for the second test. During the tests, both CSs were presented 4 times each for 20 sec with 60 sec intervals on an alternated schedule with no reward being delivered. One resting day was given in between tests.

### Behavioural measures

For both the Pavlovian conditioning and extinction tests, the measurement of the conditioned response elicited by a conditioned stimulus (CS) was determined by quantifying magazine visits during the 20 seconds of the CS, while subtracting the visits during the PreCS period, which refers to the 20 seconds preceding said CS presentation. This measure is referred to as CS-PreCS magazine visits. CS-PreCS during the first Non-degraded and Degraded CS was used as the critical measure for the Pavlovian extinction tests quantifying the cue-evoked reward-seeking response for each CS presentation. During the degradation phase, we assessed the conditioned response by dividing the magazine visits during the CS by the mean visits for that CS in the last session of conditioning (% baseline) to avoid potential contamination of the PreCS period by non-contingent reward delivery.

### Histology

At the end of behavioural experiments, rats were killed with an overdose of sodium pentobarbital (Exagon® Euthasol) and perfused transcardially with 60 ml of saline followed by 260 ml of 4% paraformaldehyde (PFA) in 0.1 M phosphate buffer (PB). Brains were removed and post-fixed in the same PFA 4% solution overnight and then transferred to a 0.1 M PB solution. Subsequently, 40 µm coronal sections were cut using a VT1200S Vibratome (Leica Microsystems). Every fourth section was collected to form a series. Immunofluorescence staining was performed for mCherry (AAV8-CaMKII-hM4D(Gi)-mCherry) and FsRed (CAV-2-PRS-FsRed) proteins: free-floating sections were first rinsed in 0.1 M phosphate buffer saline (PBS) (4×5 min) and then transferred to PBS containing 0.3% Triton X-100 (PBST) for further rinsing (3×5 min), before being incubated in blocking solution (PBST containing 3% goat serum) for 1 hr at room temperature; sections were then incubated for 24 h at room temperature with a primary antibody (rabbit polyclonal anti-RFP, 1/1000) diluted in blocking solution; after washes (4×5 min) in PBST, sections were incubated for 2 h at room temperature with the secondary antibody (goat polyclonal anti-rabbit TRITC-conjugated, 1/200) diluted in PBST; following rinses in PBS (4×5 min), sections were collected on gelatin-coated slides using 0.05 M PB, before being mounted and cover-slipped using Fluoroshield with DAPI mounting medium. DAB staining was performed for mCherry to confirm CAV-2-PRS-HA-hM4Di-hSyn-mCherry injection sites: free-floating sections were rinsed in PBST (4×5 min) before being incubated in an H2O2 (0.5%) in PBST solution for 30 min at room temperature; sections were rinsed again in PBST (4×5 min) before being incubated in the blocking solution; following blocking, they were incubated overnight at room temperature with the primary antibody (rabbit polyclonal anti-RFP, 1/2500) in blocking solution; after washing in PBST (4×5 min), sections were placed with the secondary antibody (biotinylated goat anti-rabbit, 1/1000) diluted in PBST for 2 h at room temperature; following washes (4×5 min) in PBST, they were then incubated with the avidin-biotin-peroxydase complex (1/500 in PBST) for 90 min in a dark environment at room temperature. For the final DAB staining, H202 was added to the solution (10 mg tablet dissolved in 50 mL of 0.1 M Tris buffer) just before incubation of 3-4 min; sections were then rinsed with 0.05 M Tris buffer (2×5 min) and 0.05 M PB (2×5 min), before mounting in 0.05 M PB and cover-slipping using a Fluoroshield medium.

### Tracing analysis

To quantify neuronal density within the LC, each animal received a unique identification number to allow blind counting. eGFP+ and FsRed+ neurons were captured through a Leica VM5500B Fluorescence Motorized Microscope at 10X magnification. Images were processed using MicroManager. ImageJ was used to quantify cell bodies in each section. The Paxinos rat atlas served as a reference to identify antero-posterior levels (“The Rat Brain in Stereotaxic Coordinates: Hard Cover Edition - George Paxinos, Charles Watson - Google Books,” n.d.).

### Statistical analysis

Each rat was assigned a unique identification number that was used to conduct blind testing and data analysis. A 3-way repeated measures analysis of variance (ANOVA) with Treatment (Veh vs DCZ, to be administered during the degradation phase) as a between-subjects factor and Sessions and Stimulus (Non-degraded vs Degraded CS during the degradation phase) as within-subject factors was used to analyze Pavlovian conditioning. A 3-way factorial ANOVA with Treatment at degradation (Veh vs DCZ) as a between-subjects factor and Stimulus (Degraded vs Non-degraded) and Sessions as within-subject factors was used to analyze Pavlovian degradation. A 3-way factorial ANOVA with Treatment at degradation (Veh vs DCZ) as between-subjects factors and Treatment at test (Veh vs DCZ) and Stimulus (Degraded vs Non-degraded) as a within-subject factor was used to analyze extinction tests. The accepted value for significance was p<0.05. Following significant interaction effects, the Bonferroni post-hoc test was used for multiple comparisons. Statistical analyses were performed using GraphPad Prism. Data graphs were created using GraphPad Prism and Adobe Illustrator.

## Results

### Experiment 1

In a first set of experiments, we sought to explore the potential role of mPFC and vlOFC in updating S-O associations. To this end, we implemented a Pavlovian contingency degradation task where animals had to learn that a cue, which had previously solely predicted a specific outcome, was no longer a reliable predictor. The task design is shown in **Table 1**. To investigate the involvement of both regions of interest, we used a chemogenetic strategy to selectively inhibit CaMKII+ neurons during the degradation phase, ensuring that the conditioning process remained unaltered. Rats were bilaterally injected with a AAV8-CaMKII-hM4D(Gi)-mCherry virus targeting the mPFC or the vlOFC. Then, following the 3-week recovery period, they underwent 8 daily sessions of Pavlovian conditioning during which they received two specific and counterbalanced S-O associations (15 presentations of each CS/session). Following conditioning, animals were split into two groups with similar performances. Then, they underwent 6 Pavlovian degradation sessions. 45 min before each degradation session, half of the animals received an injection of Veh, half an injection of DCZ. During each session, the predictability of one of the two cues was degraded by the non-contingent delivery of the corresponding reward during the inter-trial interval period with the frequency of delivery matching the one of the stimulus presentation, accompanied by a reduction in the likelihood of delivery at the cue itself. After completing the degradation phase, rats underwent 2 tests in extinction during which they received either Veh or DCZ, administered in a counterbalanced order. This within-subject experimental designed allowed us to assess the specific impact of chemogenetic inhibition during degradation (DCZ/Veh group), test (Veh/DCZ), or both (DCZ/DCZ), as compared to the control group (Veh/Veh). Experimental groups sizes were the following: mPFC-Veh during degradation, n=8; mPFC-DCZ during degradation, n=10; vlOFC-Veh during degradation, n=10; vlOFC-DCZ during degradation, n=11.

**Table 1.** Pavlovian contingency degradation task. At first, animals are trained over 8 daily conditioning sessions to learn that a specific CS (tone/clicker) is linked to the delivery of a specific reward (grain/sugar pellets) with a probability of 100% (p=1). Then, during 6 daily session of contingency degradation, we manipulate one of the two CSs so that the likelihood of receiving the related reward is reduced to 50%, i.e. the probability of receiving the reward is equal during (p=0.5) or outside the CS (p=0.5). At last, animals are tested in extinction (i.e. no reward being delivered) and, under normal conditions, should display a higher response to non-degraded (CS1) vs degraded cues (CS2), as measured in number of magazine visits.

| CS | Conditioning | Contingency degradation | Test |
| --- | --- | --- | --- |
| Non degraded | CS1 → US1 (100%) | CS1 → US1 (100%) | CS1 → ∅ |
| Degraded | CS2 → US2 (100%) | CS2 → US2 (50%)<br>No CS → US2 (50%) | CS2 → ∅ |

#### Histology

Figure 1A and Figure 2A show the location of AAV8-CaMKII-hM4Di-mCherry injections (with a representative image and a viral spread map) in the mPFC and vlOFC, respectively. Two rats from the mPFC group were excluded due to one-sided transgene expression.

**Figure 1.**
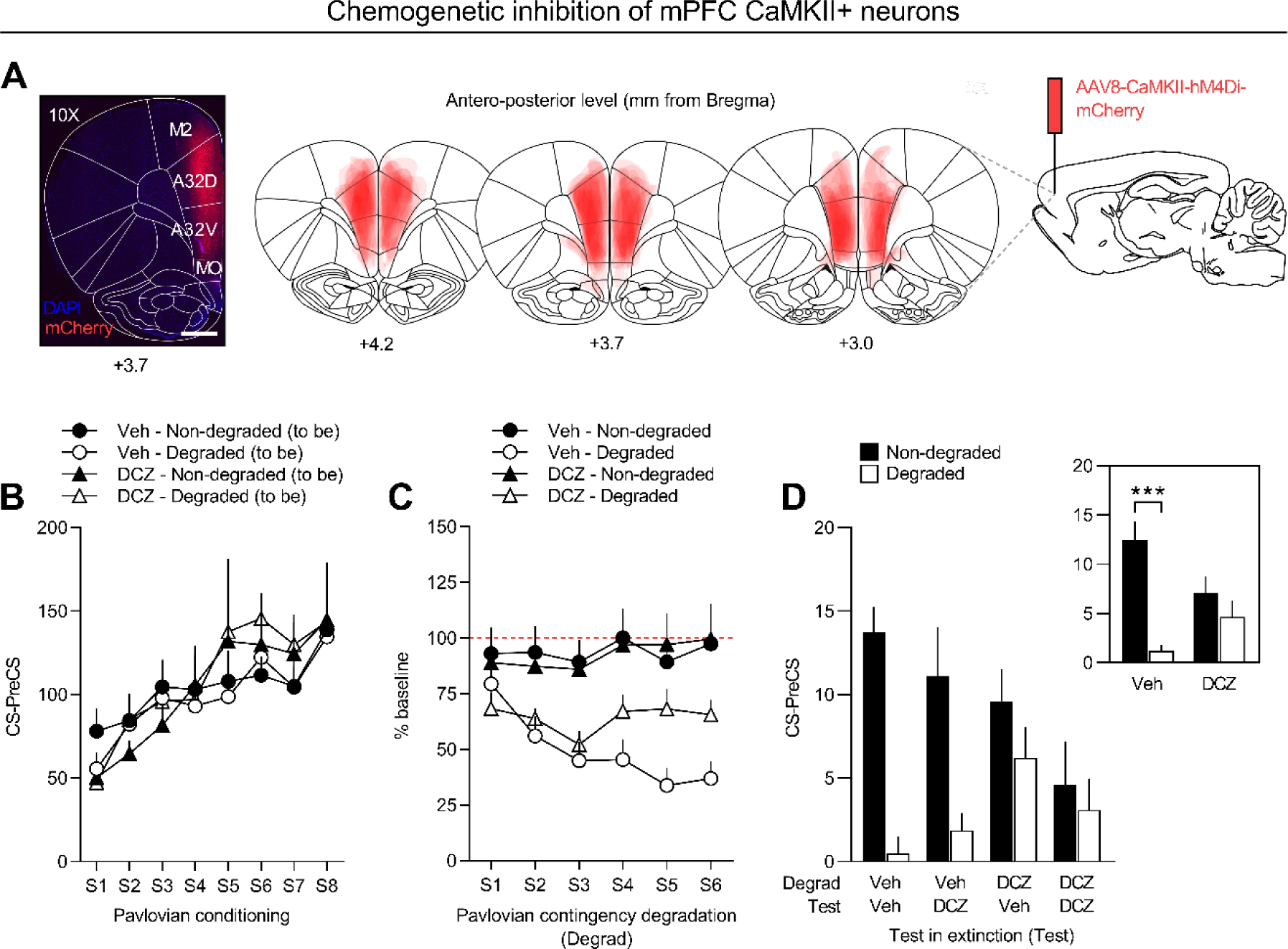
Silencing mPFC CaMKII+ neurons impairs contingency degradation learning and adaptive responding to Pavlovian S-O associations following contingency degradation. (**A**) Representative image depicting the distribution of AAV8-CaMKII-hM4Di-mCherry expression within the mPFC (A32D+A32V) and schematic representation delineating injection site and viral expression extent within the subregion for all subjects, where each rat is represented as a unique stacked layer (n=18). (**B**) Rate of CS-PreCS magazine visits across Pavlovian conditioning sessions. Data are shown based on the treatment (Veh, n=8 vs DCZ, n=10) and cue degradation (Non-degraded vs Degraded) rats will receive in the following phase. (**C**) Magazine visit rate during the CS period expressed relative to the CS magazine visits in the last session of conditioning (% baseline). Data are shown based on the treatment (Veh vs DCZ) and degradation (Non-degraded vs Degraded). (**D**) Rate of CS-PreCS magazine visits for the first presentation of each CS during the tests in extinction. Data are shown based on the treatment administered during the degradation (Degrad) and test in extinction (Test). The inlet shows data for both tests in extinction grouped for treatment administered during contingency degradation. Data are expressed as mean + SEM. ***p<0.001.

**Figure 2.**
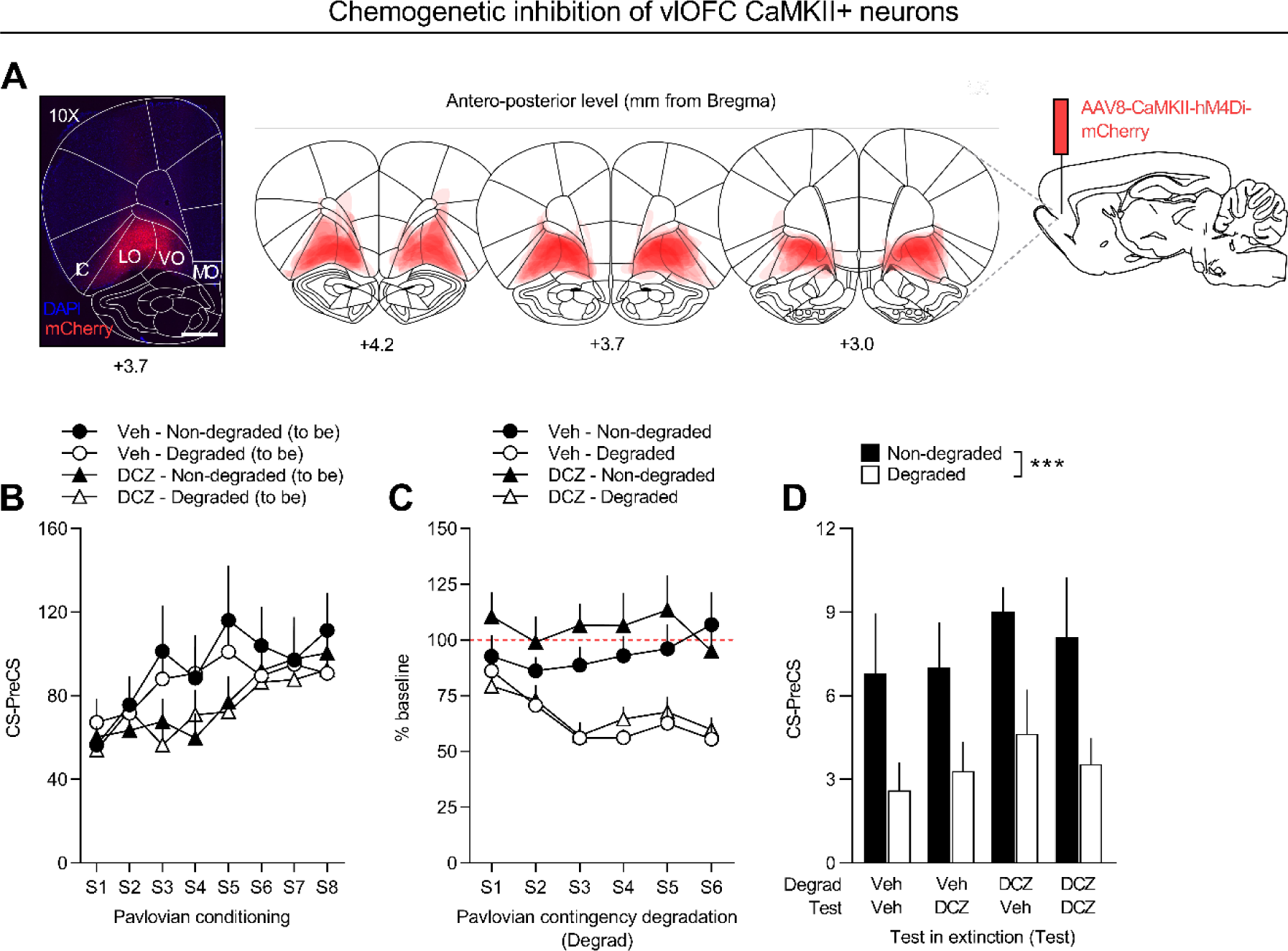
Silencing vlOFC CaMKII+ neurons does not affect adaptive responding to Pavlovian S-O associations following contingency degradation. Representative image depicting the distribution of AAV8-CaMKII-hM4Di-mCherry expression within the vlOFC (VO+LO) and schematic representation delineating injection site and viral expression extent within the subregion for all subjects, where each rat is represented as a unique stacked layer (n=21). (**B**) Rate of CS-PreCS magazine visits across Pavlovian conditioning sessions. Data are shown based on the treatment (Veh, n=10 vs DCZ, n=11) and cue degradation (Non-degraded vs Degraded) rats will receive in the following phase. (**C**) Magazine visit rate during the CS period expressed relative to the CS magazine visits in the last session of conditioning (% baseline). Data are shown based on the treatment (Veh vs DCZ) and degradation (Non-degraded vs Degraded). (**D**) Rate of CS-PreCS magazine visits for the first presentation of each CS during the tests in extinction. Data are shown based on the treatment administered during the degradation (Degrad) and test in extinction (Test). The inlet shows data for both tests in extinction grouped for treatment administered during contingency degradation. Data are expressed as mean + SEM. ***p<0.001.

#### Pavlovian conditioning

Pavlovian conditioning (Figure 1B and Figure 2B) was measured by comparing magazine visit rates during the CS (20 sec) and Pre-CS periods (20 sec prior). Stimulus-evoked response was revealed by an increased magazine visit rate during the CS period (Period) for both mPFC (F(1,32)=62.67, p<0.0001) and vlOFC animals (F(1,40)=89.80, p<0.0001). This gradual increase in responding to the CS was revealed by a significant interaction between Period and Session for both conditions (F(7,224)=2.294, p=0.0282 for mPFC and F(7,280)=2.084, p=0.0454 for vlOFC). The lack of effect of Stimulus (F(1,32)=0.2775, p=0.6020 for mPFC and F(1,40)=0.0256, p=0.8737 for vlOFC) or any interaction with this factor (F’s<1, p>0.05 for both groups) indicates that CSs were properly counterbalanced for the degradation phase and conditioning was similar for both groups. One subject failed to acquire conditioned responding to both CSs and was therefore excluded from the data analysis.

#### Pavlovian contingency degradation

Figure 1C and Figure 2C show the rate of magazine visits during each CS, relative to its corresponding baseline during the last session of conditioning, for mPFC and vlOFC rats, respectively. This measure was favored here over CS-PreCS magazine visits, given that non-contingent reward deliveries can occur during the PreCS period for the degraded CS, introducing a potential bias to this measurement. For the mPFC group (Figure 1C), both Veh- and DCZ-treated animals displayed adaptive responding, characterized by a reduced cue-evoked response to the degraded CS, while maintaining a similar response rate to the non-degraded one. This was supported by statistical analysis, which indicated a significant effect of Stimulus (F(1,16)=23.91, p=0.0002) and a significant interaction between Stimulus and Session (F(5,80)=5.298, p=0.0003). Interestingly, statistical analysis revealed a significant interaction between Session and Treatment at degradation (F(5,80)=2.835, p=0.0208). This interaction reflects an overall higher responding in DCZ-treated rats, as compared to Veh-treated controls, an effect that is driven by the degraded cue on the last 2 days of contingency degradation (p=0.0659 and p=0.0778), as confirmed by a 2-way ANOVA run exclusively on the degraded data (F(5,80)=6.168, p<0.0001). However, no significant interactions between Stimulus and Treatment at degradation (F(1,16)=1.140, p=0.3015), nor between all 3 factors (F(5,80)=1.914, p=0.1011), were detected. On the other hand, both Veh- and DCZ-treated animals of the vlOFC group displayed adaptive responding to the degraded CS (Figure 2C). This was corroborated by statistical analyses, showing a significant effect of Stimulus (F(1,19)=27.38, p<0.0001), as well as a significant interaction between Stimulus and Session (F(5,95)=5.024, p=0.0004). Overall, chemogenetic inhibition of mPFC, but not of vlOFC, CaMKII+ neurons during the Pavlovian contingency degradation impaired adaptive learning, as indicated by a higher responding to the degraded cue in comparison to controls.

#### Test in extinction

Figure 1D and Figure 2D depict the rate of magazine visits during the first presentation of each CS, subtracting the magazine visits rate during the corresponding PreCS period serving as a baseline. For the mPFC group (Figure 1D), rats treated with Veh during the degradation phase consistently exhibited a strong preference for the non-degraded conditioned stimulus (Non-degraded CS), regardless of the treatment administered during the subsequent test in extinction (Veh/Veh and Veh/DCZ). In contrast, rats treated with DCZ during the degradation phase demonstrated a comparable rate of responding for both cues, irrespective of the treatment received during tests (DCZ/Veh and DCZ/DCZ). This was confirmed by statistical analyses revealing a significant interaction between Stimulus and Treatment at degradation (F(1,16)=11.80, p=0.0034), with post-hoc tests confirming significance for Veh-treated (p<0.0001) and non-significance for DCZ-treated animals (p=0.3414). This indicates that inhibiting mPFC CaMKII+ neurons during the degradation phase impaired rats’ ability to adaptively respond to S-O associations following degradation. Moreover, the response rate during the test appeared to be lower for rats treated with DCZ, a trend that was nearly discernible in the statistical analysis, indicating a marginally non-significant effect of Treatment at test (F(1,16)=4.316, p=0.0542). No significant effects were observed for any other factors or interactions (F’s<3, p>0.05), except for an overall Stimulus effect (F(1,16)=28.60, p<0.0001). In the case of the vlOFC (Figure 2D), all groups seemed to favor the non-degraded CS, regardless of the treatment administered during the degradation phase or at the time of testing. This observation was corroborated by statistical analyses revealing a significant effect of Stimulus (F(1,19)=13.43, p=0.0016), while all other factors yielded non-significant results (F’s<2, p>0.05). Overall, chemogenetic inhibition of mPFC, but not of vlOFC, CaMKII+ neurons during the Pavlovian contingency degradation impaired adaptive responsive in unrewarded testing conditions, independently from treatment at testing, as indicated by a similar responding to non-degraded and degraded cues.

### Experiment 2

In light of the differential effects observed in Experiment 1 and the hypothesized influence of the LC-NA system in prefrontal-dependent learning processes, as a next step we aimed at characterizing NA projections to the mPFC and the vlOFC. To do so, we employed a unilateral double-retrograde viral strategy to simultaneously target the mPFC and the vlOFC in the same individuals (n=8). One region was injected with a green CAV-2 vector (CAV-2-PRS-eGFP), while the other was injected with a red CAV-2 vector (CAV-2-PRS-FsRed), counterbalancing hemispheres and regions between animals. These vectors allowed the expression of either fluorescent protein under the control of a synthetic DβH promoter (PRS), therefore enabling selective targeting of NA neurons projecting to the subregions of interest. This allowed us to count the number of eGFP- and FsRed-positive cell bodies within the LC, i.e. quantify the amount of NA neurons projecting to either one subregion, the other, or both (Figure 3A). Quantification of neuronal density within the LC region revealed 37% of labelled cell bodies projecting to the mPFC (517/1403 labelled somas), 49% projecting to the vlOFC (689/1403 labelled somas), and 14% projecting to both (197/1403 labelled somas), suggesting relatively segregated neuronal populations (Figure 3B). Among cells projecting to the mPFC, 28% also projected to the vlOFC, while 21% of vlOFC-projecting cells concurrently projected to the mPFC. Importantly, we found no difference of expression between neither vectors nor hemispheres. Next, we examined the distribution of labelled cell bodies within the LC along its antero-posterior axis (Figure 3C). Our analysis revealed no distinct topographical organization along either the antero-posterior or dorso-ventral axis. Instead, the organization of both mPFC and vlOFC-projecting neurons appeared to align with the surface of the LC at each antero-posterior level, with no observed differences between those projecting to either region or both (Figure 3D). Overall, the significant degree of segregation observed among NA projections to the mPFC and the vlOFC prompted us to investigate whether inhibiting these projections specifically would also result in a subregion-specific deficit and therefore offer a neuromodulatory interpretation for what observed in Experiment 1.

**Figure 3.**
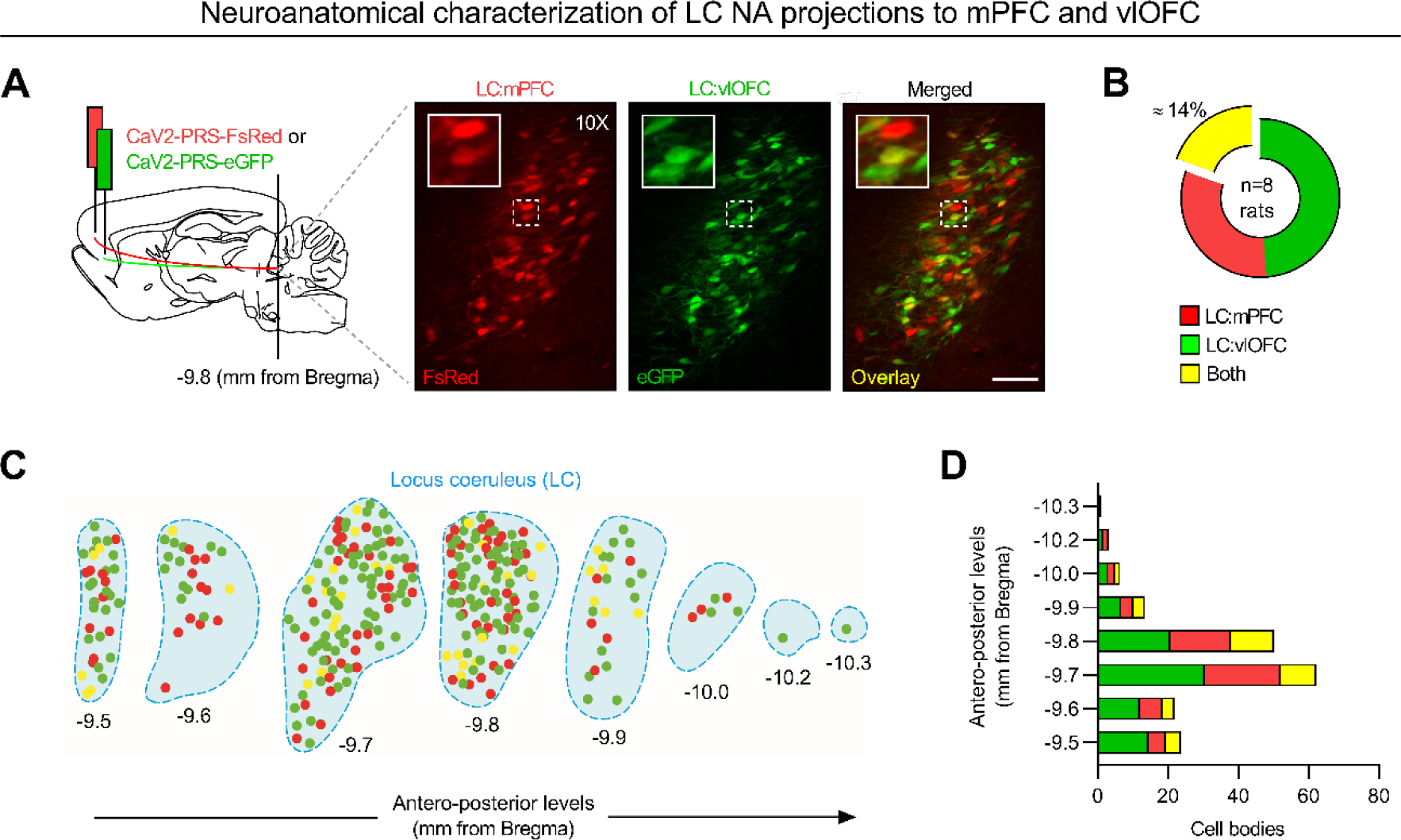
LC NA cells exhibit target-specific projections. (**A**) Schematic illustrating unilateral double-viral injections of retrograde CAV-2-PRS-eGFP and CAV-2-PRS-FsRed vectors within mPFC and vlOFC and representative immunofluorescent staining images of corresponding FsRed+, eGFP+ and overlay retrogradely traced NA cell bodies within the LC. Scale bar: 100 µm. (**B**) Overall quantification of NA cell bodies projecting to vlOFC (green), mPFC (red) or both (yellow) across all rats (n=8). (**C**) Spatial organization of noradrenergic cell bodies projecting to vlOFC (green) mPFC (red) or both (yellow) along the antero-posterior axis (from −9.5mm to −10.3mm relative to Bregma) in two representative rats (one injected in the left, the other in the right hemisphere). (**D**) Spatial distribution of NA cell bodies projecting to vlOFC (green), mPFC (red) or both (yellow) along the LC antero-posterior axis across all rats (n=8). Data are expressed as sum of cell bodies.

### Experiment 3

Considering the observed impact of manipulating mPFC and vlOFC neuronal activity on S-O updating, along with the substantial segregation of NA projections to these regions, in a last set of experiments we aimed to explore whether chemogenetic inhibition of these sets of projections would yield any effect on S-O updating. To this end, rats were bilaterally injected with a retrograde CAV-2-PRS-HA-hM4Di(Gi)-hSyn-mCherry vector within the designated mPFC and vlOFC regions, in order to enable selective chemogenetic targeting of NA projections. Subsequently, the animals underwent the same recovery and behavioural procedures as outlined in Experiment 1. Experimental groups sizes were the following: mPFC-Veh during degradation, n=9; mPFC-DCZ during degradation, n=10; vlOFC-Veh during degradation, n=13; vlOFC-DCZ during degradation, n=13.

#### Histology

Figure 4A and Figure 5A illustrate the spread of CAV-2-PRS-HA-hM4Di(Gi)-hSyn-mCherry injections in the mPFC and vlOFC, respectively, together with a representative picture of mCherry-expressing retrogradely targeted neurons in the LC. We previously showed (Cerpa et al., 2023) that while mCherry staining is also present at injection sites, reflecting local cortico-cortical connections that are not NA dependent, HA-immunoreactive cell bodies were found exclusively in the LC, observations consistent with NA-specific expression of the HA-tagged hM4Di due to the PRS promoter, and nonselective expression of mCherry due to the hSyn promoter. However, considering that mCherry and HA immunoreactivity fully colocalize in the LC, in the current study we opted to use mCherry as a proxy of DREADDs expression.

**Figure 4.**
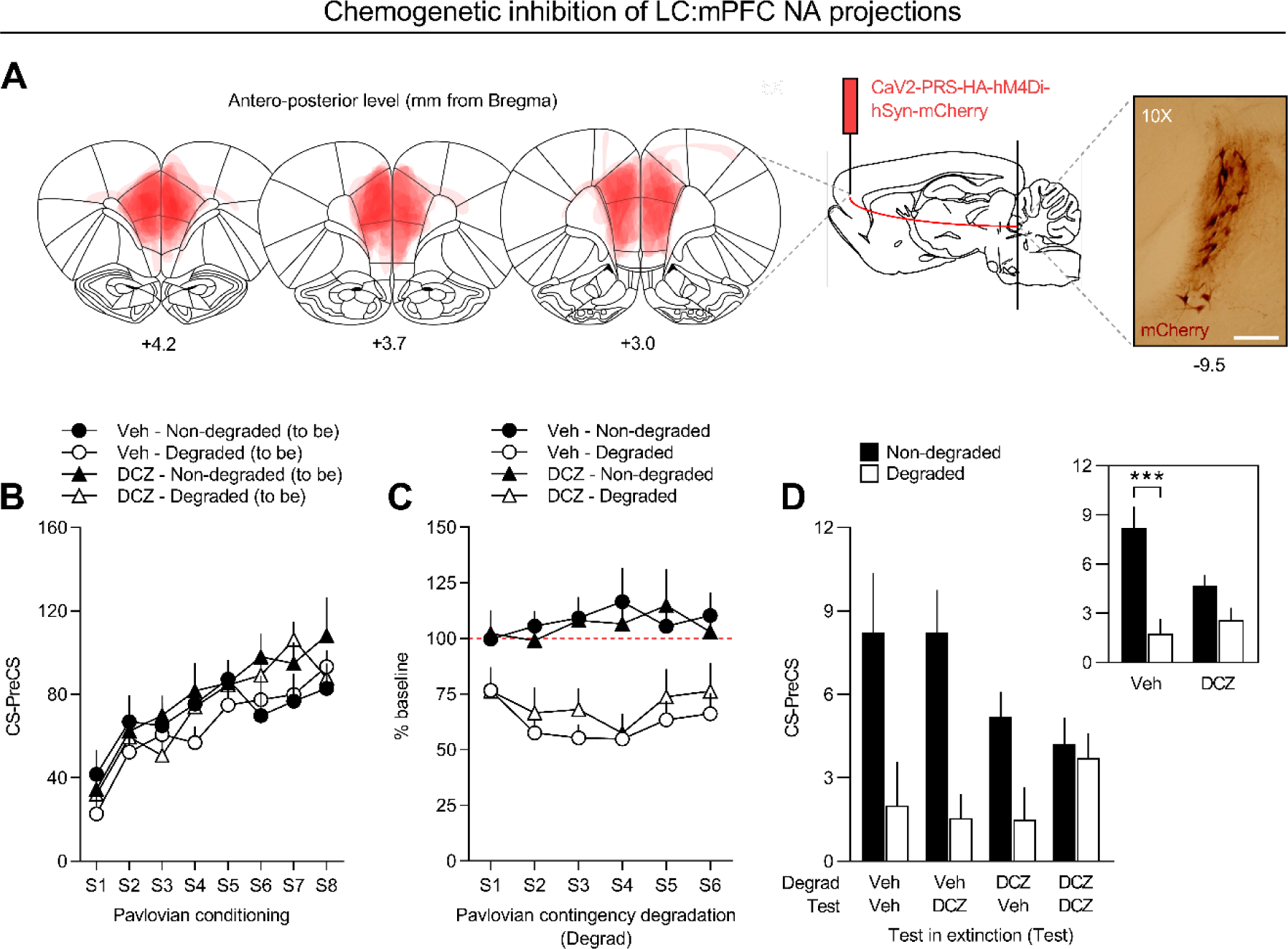
Silencing mPFC NA inputs impairs adaptive responding to Pavlovian S-O associations following contingency degradation. (**A**) Schematic representation delineating viral expression extent within the mPFC (A32D+A32V) for all subjects, where each rat is represented as a unique stacked layer (n=19), and representative image showing retrogradely targeted LC cell bodies. Scale bar: 50 µm. (**B**) Rate of CS-PreCS magazine visits across Pavlovian conditioning sessions. Data are shown based on the treatment (Veh, n=9 vs DCZ, n=10) and cue degradation (Non-degraded vs Degraded) rats will receive in the following phase. (**C**) Magazine visit rate during the CS period expressed relative to the CS magazine visits in the last session of conditioning (% baseline). Data are shown based on the treatment (Veh vs DCZ) and degradation (Non-degraded vs Degraded). (**D**) Rate of CS-PreCS magazine visits for the first presentation of each CS during the tests in extinction. Data are shown based on the treatment administered during the degradation (Degrad) and test in extinction (Test). The inlet shows data for both tests in extinction grouped for treatment administered during contingency degradation. Data are expressed as mean + SEM. ***p<0.001.

**Figure 5.**
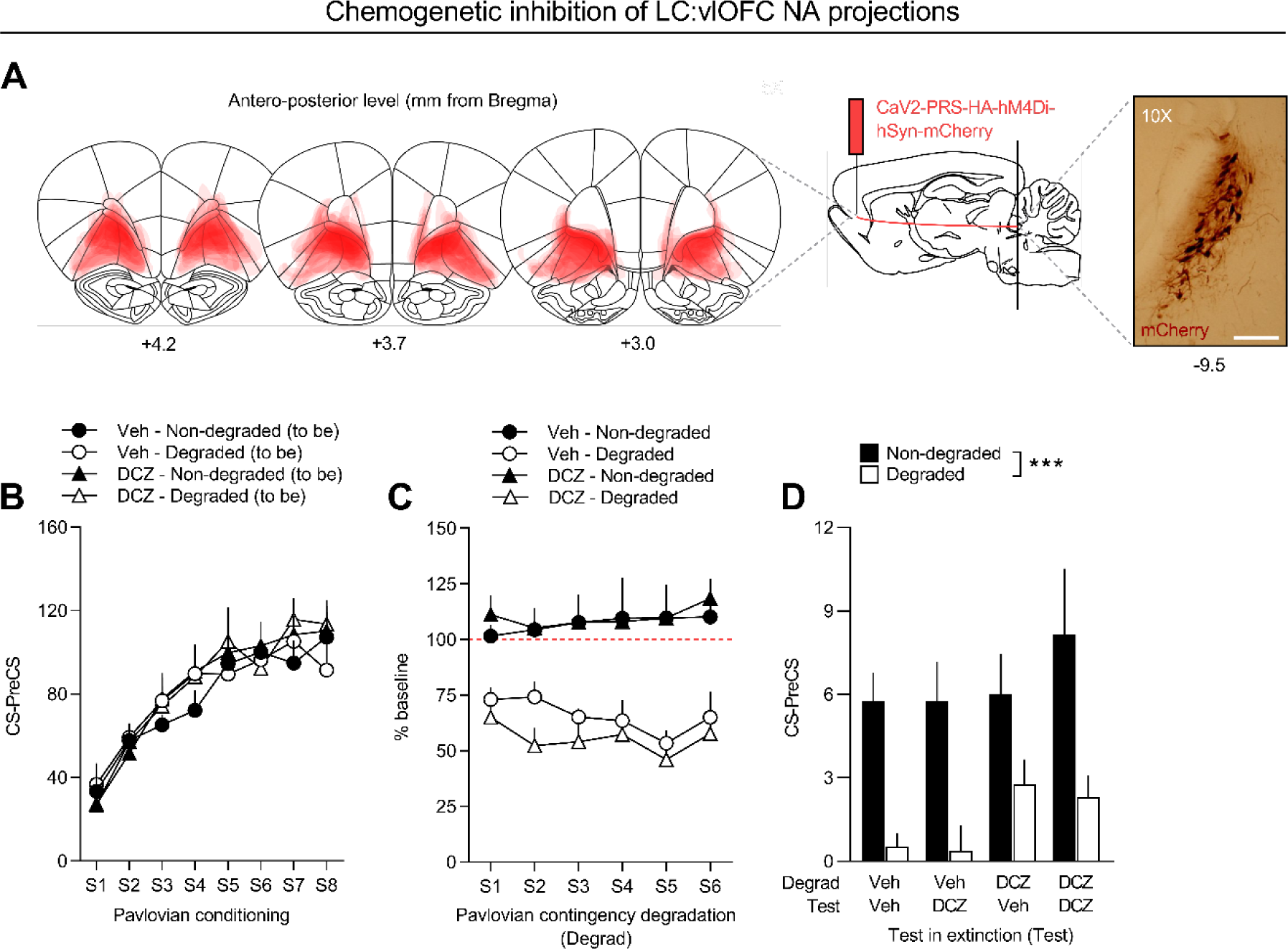
Silencing vlOFC NA inputs does not affect adaptive responding to Pavlovian S-O associations following contingency degradation. (**A**) Schematic representation delineating viral expression extent within the vlOFC (VO+LO) for all subjects, where each rat is represented as a unique stacked layer (n=19), and representative image showing retrogradely targeted LC cell bodies. Scale bar: 50 µm. (**B**) Rate of CS-PreCS magazine visits across Pavlovian conditioning sessions. Data are shown based on the treatment (Veh, n=13 vs DCZ, n=13) and cue degradation (Non-degraded vs Degraded) rats will receive in the following phase. (**C**) Magazine visit rate during the CS period expressed relative to the CS magazine visits in the last session of conditioning (% baseline). Data are shown based on the treatment (Veh vs DCZ) and degradation (Non-degraded vs Degraded). (**D**) Rate of CS-PreCS magazine visits for the first presentation of each CS during the tests in extinction. Data are shown based on the treatment administered during the degradation (Degrad) and test in extinction (Test). Data are expressed as mean + SEM. ***p<0.001.

#### Pavlovian conditioning

Pavlovian conditioning (Figure 4B and Figure 5B) was assessed by comparing magazine visit rates during the CS (20s) and Pre-CS periods (20s prior). The stimulus-evoked response was evident through an elevated magazine visit rate during the CS period (Period) for both mPFC (F(1,36)=222.8, p<0.0001) and vlOFC animals (F(1,50)=131.8, p<0.0001). The progressive increase in CS-evoked response was evident through a significant interaction between Period and Session for both conditions (F(7,252)=6.955, p<0.0001 for mPFC and F(7,350)=22.13, p<0.0001 for vlOFC). No effect of Stimulus or any interaction with this factor were observed for the vlOFC group (F’s<2, p>0.05). However, despite the absence of interactions for the mPFC group (F’s<2, p>0.05), Choice was found to be statistically significant (F(1,36)=5.508, p=0.0245). However, no effect was found for Choice when analyzing a CS-PreCS measure (F(1,17)=2.312, p=0.1468) nor an interaction between Choice x Treatment (F(1,17)=0.069, p=0.796).

#### Pavlovian contingency degradation

Similar to our first experiment, we opted for the last session of conditioning as a baseline for responding to each CS, rather than relying on the corresponding Pre-CS period. We observed a similar impact of degradation in mPFC and vlOFC groups, with evidence of adaptive responding, characterized by a reduction in the CS-evoked response to the degraded cue, while the response to the non-degraded CS remained unchanged compared to their respective baselines (Figure 4C and Figure 5C). This was substantiated by statistical analysis, indicating a significant effect of Degradation (F(1,17)=20.74, p=0.0003 for mPFC and F(1,24)=46.71, p<0.0001 for vlOFC) and an interaction between Degradation x Session (F(5,85)=2.835, p=0.0204 for mPFC and F(5,120)=2.937, p=0.0155 for vlOFC). Overall, this suggests that both the mPFC and vlOFC groups appeared to have successfully adapted to the degraded cue, irrespective of the treatment received during the degradation phase.

#### Test in extinction

In the mPFC group (Figure 4D), animals treated with DCZ during the degradation and the test phases appeared to exhibit no preference for the non-degraded cue, while Veh-treated animals during the degradation showed a preference, regardless of their treatment at test, confirming proper learning of the degradation. Indeed, statistical analysis revealed a significant interaction between Degradation x Treatment at degradation (F(1,17)=7.973, p=0.0117), with post-hoc analyses confirming that Veh-treated animals exhibited a preference for the non-degraded CS (p<0.0001), while DCZ-treated animals did not (p=0.1275). Overall, these findings indicate that NA projections to the mPFC, similarly to mPFC CaMKII+ neurons, are required for updating Pavlovian S-O associations in the context of contingency degradation. Additionally, although responding seemed lower for the DCZ-treated group during degradation, no significant of Treatment at degradation effect was found (F(1,17)=1.309, p=0.2685). No significant effects were observed for any other factors or interactions (F’s<2, p>0.05), except for an overall Degradation effect (F(1,17)=30.84, p<0.0001). Regarding the vlOFC group (Figure 5D), we observed comparable results to our initial experiment targeting CaMKII+ neurons within the same region. All groups, regardless of the treatment administered during the degradation phase or the subsequent tests, appeared to demonstrate proper learning of the degradation. This was confirmed by a significant effect of Degradation (F(1,24)=24.18, p<0.0001), with all other factors or interactions being non-significant (F’s<3, p>0.05). Overall, chemogenetic inhibition of NA projections to the mPFC, but not to the vlOFC, during the Pavlovian contingency degradation impaired adaptive responsive in unrewarded testing conditions, independently from treatment at testing, as indicated by a similar responding to non-degraded and degraded cues.

## Discussion

The significance of incoming signals frequently changes in natural settings. This solicits organisms to regularly track the ongoing predictive value of environmental cues and adjust their behavior accordingly. Degrading the contingency between a stimulus (S) and its associated outcome (O) is one effective way to study such adaptive behavior. The current study presents evidence of medial prefrontal cortex (mPFC) neurons, and more specifically of noradrenergic (NA) transmission within the subregion, being crucial for adaptive responding following Pavlovian contingency degradation.

A wealth of literature has shown that the mPFC plays a crucial role in mediating the degradation of instrumental action (A)-O contingencies in rodents (Woon et al., 2020). Indeed, neurotoxic lesions of the prelimbic subdivision of the mPFC (A32D) prevented this update in rats (Balleine and Dickinson, 1998; Corbit and Balleine, 2003; Coutureau et al., 2012), as did chemogenetic silencing in mice (Swanson et al., 2017), an effect that is believed to be dopamine-dependent, as suggested by both DA lesions and inactivation of D1/D2 receptors (Naneix et al., 2009; but see Lex and Hauber, 2010), and mediated by ventral hippocampal (vHPC) inputs, as shown by chemogenetic synaptic silencing of vHPC terminals (Piquet et al., 2023). A relatively low cross-species homology in prefrontal functioning hampered a clear-cut translation of these findings in non-human primates (Duan et al., 2021), although reports implicating subdivisions of the mPFC in instrumental degradation exist (Jackson et al., 2016). Whether the mPFC is implicated in the degradation of Pavlovian S-O contingencies, on the other hand, remained a long-standing question. In here, we show that chemogenetic inhibition of mPFC (A32D+A32V), but not of ventromedial OFC (vlOFC, VO+LO), CaMKII+ neurons impairs the rats’ ability to both learn the degradation and appropriately respond between non-degraded and degraded cues in unrewarded testing conditions. This suggests that mPFC neurons play a key role in both encoding and recalling degraded S-O associations. Worth mentioning, an important division of labor has been proposed in which the prelimbic (A32D) and infralimbic (A32V) subdivisions of the mPFC regulate the expression and suppression of adaptive behaviors (Quirk et al., 2000; Milad and Quirk, 2002; Rosenkranz et al., 2003; Killcross and Coutureau, 2003; Maren and Quirk, 2004). This division of labor remains to be investigated in the framework of Pavlovian contingency degradation, and surely represents an interesting direction for future investigations.

The fact that chemogenetic inhibition of vlOFC CaMKII+ neurons did not affect the animals’ ability to learn the degradation or discriminate between cues in unrewarded testing conditions is a result in apparent contrast with what previously reported (Ostlund and Balleine, 2007; Alcaraz et al., 2015). Different variables might account for discrepancies between the current and these previous studies. First of all, there are evident differences relative to target sites, with lesioning procedures spreading either more laterally or more anteriorly within the OFC, as compared to our viral strategy, which consistently targeted VO and LO, leaving the anterior insular, medial orbitofrontal, prelimbic and infralimbic cortices untouched. Considering that relatively close prefrontal areas – or even the same area alongside its axis – often account for different behavioral effects (Fisher et al., 2020; Alsiö et al., 2020; Cerpa et al., 2023), we cannot exclude that inconsistencies in neuroanatomical targeting might represent an important source of variability. Secondly, when examining behavioral protocols, core experimental differences arise between the current and the abovementioned studies, including the number of sessions and reward deliveries, the testing procedure (rewarded vs unrewarded conditions), and – most importantly – the degradation procedure itself. Specifically, in contrast to Ostlund and Balleine, who only manipulated the likelihood of receiving the outcome after the cue (backward transition probability) while keeping the reward probability during the cue (forward transition probability) unchanged, we modified both forward and backward probabilities. Consequently, the impairment observed in their study could be tied to exclusive modulation of the backward probability. At last, we cannot exclude that differences in the rats’ experimental baggage, in the silencing techniques (post-conditioning lesions vs DREADD-mediated inhibition), as well as in the behavioral measures adopted (entries/min vs entries as a % of baseline responding), might also represent a potential source of variability.

A critical aspect of the neuronal mechanisms involved in adaptive behavior involves catecholaminergic neuromodulation, which can be functionally linked to detecting novelty, producing and interpreting prediction errors and sustaining working memory. Neurons in the locus coeruleus (LC) are believed to track uncertainty by phasically responding to novel stimuli (Bouret and Sara, 2004; Tait et al., 2007; McGaughy et al., 2008; Tervo et al., 2014; Uematsu et al., 2017; Jahn et al., 2018; Cope et al., 2019). Compelling theoretical models hypothesize that the LC interacts with the prefrontal cortex to support adaptive behaviors (Sara and Bouret, 2012; Cerpa et al., 2021). Specifically, when a change in contingencies is detected, a rise in prefrontal NA activity is believed to trigger behavioral adaptations (Aston-Jones et al., 1997; Bouret and Sara, 2005; Sadacca et al., 2017). However, these NA bursts are believed to be highly segregated, in line with the undergoing conception of the LC as a non-homogenous nucleus, in both its structure and function (Chandler et al., 2019; Poe et al., 2020).

In here, using NA-specific CAV-2-PRS constructs (del Rio et al., 2019; Hayat et al., 2020), we found that direct and discrete LC-NA projections to mPFC and vlOFC exist, thereby adding to the growing evidence of a highly modular LC architecture (Agster et al., 2013; Chandler et al., 2013, 2014, 2019). In a seminal paper, Chandler et al. already revealed largely segregated population of cells projecting to various cortical regions using retrograde tract tracing, in particular OFC vs M1, mPFC vs M1, and ACC vs M1 (Chandler et al., 2014). However, to the best of our knowledge, we are the first to directly compare mPFC vs vlOFC LC-NA projections, as well as to describe the gradient of LC-NA projections throughout the antero-posterior axis of the LC. Most importantly, we found that chemogenetic inhibition of LC-NA projections to the mPFC, but not to the vlOFC, impaired the rats’ ability to correctly respond between non-degraded and degraded cues in unrewarded testing conditions. This result suggests that NA might be specifically responsible for the distinctive behavioral outcomes observed in the first set of experiments and, more specifically, for the inability to recall degraded S-O associations in unrewarded testing conditions.

Disentangling the role of different prefrontal subregions and neurotransmitters is a key step to advance our knowledge of adaptive and maladaptive behaviors. In this sense, the current study belongs to a recent and ever-growing body of work (Cerpa et al., 2021, 2023; Chernoff et al., 2023; Cremer et al., 2023) that not only is integrating and slowly revising the classical conceptions linking the mPFC to instrumental learning and the OFC to Pavlovian learning, but suggesting a major role for NA in the neuromodulatory computations that underlie higher-order cognitive processes. For instance, a similar, but opposite, dissociation of effects was previously observed in another recent work from our laboratory (Cerpa et al., 2023). In such study, also using the CAV-2-PRS-HA-hM4Di-hSyn-mCherry vector, we showed that LC-NA projections to the vlOFC, but not to the mPFC, are required for both encoding and recalling the identity of an expected instrumental outcome, specifically when that identity has been reversed. Furthermore, another recent study, this time using a pharmacological approach, highlighted a double dissociation between the behavioral effects of NA transmission across these prefrontal subregions, this time with respect to risky choice and impulsive action (Chernoff et al., 2023).

The cellular and molecular mechanisms underlying these NA-mediated effects remain to be elucidated. However, new overarching theories of LC function, like the global model failure system theory (Jordan, 2023), postulate that cortical top-down predictive inputs to the LC, i.e. prediction errors (PE), are followed by bottom-up responsive outputs that ultimately control synaptic integration, driving behavioral change. Under this conception, novel and uncertain environments would prompt high rates of PE, resulting in high LC outputs. The current work, together with our previous study (Cerpa et al., 2023), provides behavioral evidence of a strong cortical parcellation in these bottom-up responses, in line with recent conceptions of prefrontal functioning relatively to goal-directed behavior (Turner and Parkes, 2020; Cerpa et al., 2021). We propose that, while vlOFC-NA projections might be implicated in adaptations to action violations and outcome identity changes, as happens in reversal learning, mPFC-NA projections might respond to context-dependent incentive memory violations, as for the case of instrumental and Pavlovian degradation. Whether DA transmission in the mPFC is also required for the latter form of adaptive responding remains to be tested.

In summary, the current findings provide clear evidence of prefrontal parcellation of function in the framework of adaptive responding. Our results question the previously assumed role of the OFC in Pavlovian contingency degradation, and point instead at the mPFC as a key player. Notably, we show that this effect is driven – at least in part – by NA innervation of the subregion, adding to current theories suggesting a major, but complex and modular, role for the LC in the regulation of flexible, goal-directed decision-making.

## Acknowledgements

This work was supported by the French National Research Agency (CE37-0019 NORAD to EC). The authors wish to thank Angélique Faugère for her help with immunofluorescence and Yoan Salafranque for his expert animal care.

